# Forest conversion to cashew orchards delays the diel activity of diurnal mammals in West Africa

**DOI:** 10.1101/2025.04.01.646119

**Authors:** Ana Filipa Palmeirim, Daniel Na Mone, Isnaba Nhassé, João Soares, Raquel Oliveira, Manuel Lopes-Lima, Luís Palma

**Author notes:** Corresponding author: Laboratório de Zoologia e Ecologia de Vertebrados, Instituto de Ciências Biológicas, Universidade Federal do Pará, Av. Perimetral, 2-224, 66077-830 Belém, Pará, Brasil. Teaser text Mammals in West Africa use both forests and cashew orchards, but diurnal species shift their activity later in the day, possibly to reduce encounters with humans. Nocturnal species show no changes, suggesting different responses to human-modified landscapes.

## Abstract

Land-use change is the main driver of biodiversity loss worldwide. However, less is known about how species adapt their behavior to persist in human-altered habitats. We addressed this gap by examining mammal responses in terms of habitat use and diel activity patterns to forest conversion into cashew orchards in West Africa. We surveyed mammals using camera-trapping in 12 forest sites and 12 cashew orchards in the Cantanhez National Park in southwest Guinea-Bissau, West Africa. We grouped mammals into (1) diurnal and cathemeral and (2) nocturnal species according to their diel activity periods and assessed their use and diel activity between habitat types. Based on 709 trap-days, we obtained 842 records of 12 diurnal or cathemeral species (18.3% of the records) and 13 nocturnal species (81.7%). Mammal habitat use was similar between forests and cashew orchards for both diurnal/cathemeral and nocturnal species. However, the peak of diurnal/cathemeral species shifted from the morning period (∼11h00) in the forests to the afternoon period in the cashew orchards (∼16h30), resulting in little overlap in activity of diurnal/cathemeral species between habitat types. No differences were observed in the diel activity of nocturnal mammals.

Although mammals can make use of altered habitats, diurnal species are able to delay their activity, probably to avoid direct contact with humans. Our study highlights the ability of species in adapting their behaviour to persist in newly human-modified landscapes in the tropics.

## INTRODUCTION

Tropical forests, holding over 75% of all species described to science, are under increasing pressure from agricultural expansion and intensification (IUCN 2022; Powers and Jetz 2019). In West Africa, Guinea-Bissau is experiencing an unprecedent expansion of cashew cultivation (Catarino et al. 2015), having lost 77% of its closed-canopy forests between 2001 and 2018 (UN-FCCC 2019). Today, the country’s economy is heavily dependent on cashew exports, which account for 90% of the GDP (UN-FCCC 2019). Despite the potentially far-reaching impacts of this widespread land-use change on biodiversity (Guedes et al. 2024), evidence-based knowledge is largely unavailable across the Western Afrotropics (Luiselli et al. 2019).

The impacts of land-use change on biodiversity are multifaceted, shaped by both species-specific traits (Newbold et al. 2016) and the extent of structural and compositional alterations compared to the original habitat (Barlow et al. 2016). Anthropogenic disturbances often lead to local extinctions of species that are sensitive to such changes (Palmeirim et al. 2017). However, species that persist may adapt by altering their habitat use and behaviour to survive in the altered environment (Mazza et al. 2020; Tranquillo et al. 2023). Drastically modified habitats typically exhibit pronounced differences, including shifts in resource availability, microclimatic conditions, light exposure (e.g., in newly cleared areas), human activity, altered species interactions, and increased competition (Monterroso et al. 2014).

These factors can interact to drive changes in species’ habitat preferences (Ferreira et al. 2022) and their diel activity patterns, i.e. how animals distribute their activity over the 24 h day (Ikeda et al. 2016). For instance, several species changed diel activity patterns when human observers are present at waterholes in Namibia, which may preclude changes in important prey-predator interactions that shape ecosystem functioning (Patterson et al. 2024).

Tropical mammal communities exhibit exceptional taxonomic and functional diversity, making them integral to ecosystem processes. These mammals contribute to essential services that support human well-being, such as pollination, pest control, and soil bioturbation (Lacher et al. 2019). In general, terrestrial mammals have four types of activity patterns: diurnal, nocturnal, crepuscular, and cathemeral (Bennie et al. 2014). As human land-use change intensifies, wild mammal populations typically decline (Ceballos et al. 2017) and species assemblages undergo significant shifts in composition (Foord et al., 2018). Following anthropogenic disturbance, mammals are known to adjust their diel activity (Gallo et al. 2022). For example, some mammals modify their diel activity patterns to avoid human activity (Ikeda et al. 2022) or, alternatively, to take advantage of differences in temporal resource availability, as in the case of insectivorous bats on the Afrotropical Island of São Tomé (Araújo-Fernandes et al. 2025). In addition, given that human activity is generally diurnal, in human-modified environments, diurnal mammal species may be more affected than nocturnal species, in which case human disturbance can lead to “temporal” habitat loss (Rivera et al. 2022). Furthermore, diurnal species have been found to increase their nocturnal activity in urbanised areas (Carter et al. 2012), presumably to avoid periods when humans are most active (Gaynor et al. 2018a, b).

Here, we aimed to investigate diel activity responses to forest conversion into cashew orchards of mid-sized mammals in West Africa. We surveyed mammals using camera-trapping at 24 sampling sites, with 12 being in forests and 12 in cashew orchards, in the Cantanhez National Park, Guinea-Bissau. We hypothesised that (1) mammal habitat use, particularly of diurnal species, is lower in cashew orchards than in forests, and that (2) mammal activity patterns in the cashew orchards, particularly of diurnal species, might shift to avoid human contact, resulting in lower overlap of activity between habitat types.

## METHODS

*Study area.—*This study was conducted in the 1,057 km² Cantanhez National Park (CNP), designated in 2008 in the Tombali region of Guinea-Bissau (10°55’–12°45’ N, 13°37’– 16°43’ W; Fig. 1a). The park encompasses a mosaic of habitats, including sub-humid closed-canopy forests, mangroves, wet grasslands, cashew orchards, agricultural fields, and small human settlements (Pereira et al. 2022). The climate is humid monsoonal with a clear distinction between the dry season (November–May) and the wet season (June–October).

**Figure 1.**
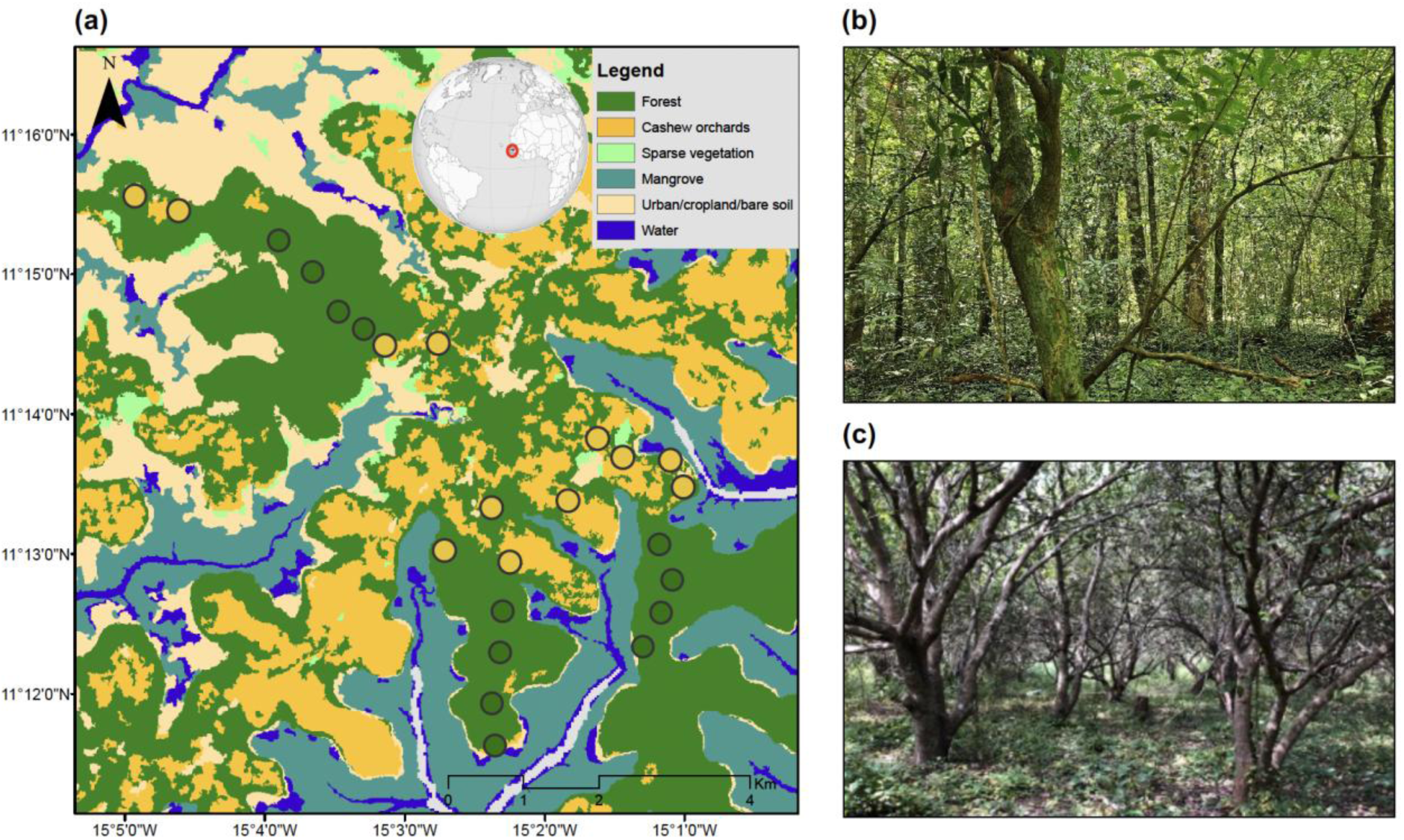
Location of the sampling sites within the study area. (a) Map of the Cantanhez National Park coloured according to the different types of land-use as extracted from Pereira et al. (2022). The inset shows the study area in detail, including the sampling sites located in the forest (green circles) and in the cashew orchards (orange circles). The sampled land-use types are further illustrated by a picture: (b) forest, and (c) cashew orchard. Photo credits: A.F. Palmeirim.

Average temperatures range from 28°C to 31°C, and the area receives an annual rainfall of roughly 2,400 mm (Climate Knowledge Portal 2021).

Although highly fragmented by agriculture and human settlements, the Park still contains forest patches that serve as important wildlife corridors (Bersacola et al. 2022). The agricultural landscape is dominated by cashew orchards, which consist of a monoculture of *Anacardium occidentale* L., a species native to Brazil. Cashew was introduced and promoted in Guinea-Bissau by the colonial administration since the early 20^th^ century (Temudo and Abrantes 2014). Cashew orchards are typically small-scale, locally managed, and cultivated without the use of agrochemicals or irrigation; trees are planted 4 to 5 metres apart, with the harvest season running from March to June. From October to December, the trees remain fruitless, and the understorey vegetation varies from sparse to moderate. In December, farmers clear the orchards, removing undergrowth to prepare for the next growing cycle (Sierra-Baquero et al. 2024).

*Mammal surveys.—*We carried out the mammal surveys between November and December 2023 using camera traps. A single digital camera (Browning Patriot model BTC-PATRIOT-FHD) was deployed at each sampling site and operated continuously for 30 days. Sampling sites were located within a landscape that included a mosaic of forest and cashew habitats (Fig. 1b-c). Cameras were strategically positioned to maximise mammal detection while minimising disturbance from human activity, namely by avoiding frequently used trails wherever possible. Site selection was guided by local nature guide (Mamadu Cassamá). Each camera was programmed to take a sequence of five photos with 15-second intervals between consecutive sequences. Cameras were unbaited, mounted on tree trunks 30–40 cm above the ground, and positioned at least 300 metres apart to maximise independence of observations.

The photographs collected during the study were analysed using TimeLapse Software (Greenberg et al. 2019; Greenberg 2023). Each image was carefully observed to identify mammals to the species level whenever possible. Consecutive images of the same species taken within a 30-minute interval were grouped as a single detection event (i.e., one record) (Gessner et al. 2014) unless individuals could be differentiated by age, sex, or distinctive physical features. Of the total 858 recorded images, 16 (1.9%) were excluded from subsequent analysis due to poor image quality or incomplete captures that prevented accurate species identification. Differences in the number of trapping days per site, which ranged from 15 to 30, were caused by logistical constraints. To account for this, detection records were standardised to a 10-day trapping period. The total sampling effort across all sites corresponded to 709 camera-trap days (Table S1). Fieldwork was conducted with the permission of the Instituto da Biodiversidade e das Áreas Protegidas (IBAP) of Guinea-Bissau, and consent was obtained from all cashew farm owners of the study area.

*Data analysis.—*Species diel activity guilds were retrieved from Wilman et al. (2014). As strictly diurnal species accounted for only 12.5% of the records (*N* = 105 records from seven species), we considered together species with (1) diurnal activity, (2) both diurnal and crepuscular activity, and (3) both diurnal, crepuscular, and nocturnal activity (i.e., cathemeral species) (Table S2). For convenience, these three categories of species activity patterns are hereafter referred to as “diurnal”. To ensure data representativeness in the analysis, species were categorised into two activity guilds—diurnal and nocturnal—and analysed accordingly.

To investigate whether the habitat use of diurnal or nocturnal species (given by the standardised number of records) was affected by habitat type (forest or cashew orchard), we performed Generalised Linear Models (GLMs). The number of records was log-transformed to achieve data normality and the models were fitted with a Gaussian distribution. Model residuals were then examined using the ‘dharma’ R package (Hartig 2022), with no issues being reported.

Diel activity patterns were first assessed by grouping all species into two activity guilds—diurnal and nocturnal, as before. Additionally, the four most frequently recorded species (i.e., those with more than 20 detections per habitat type) were analysed individually in species-specific analysis (Meredith and Ridout 2014). Time data were first transformed into radians, ranging from zero hours (0) to 24 hours (2π). Kernel density functions were then used to visualise activity patterns and calculate overlap coefficients. A smoothing parameter (kmax) of three was applied for Kernel density estimation, a value known for providing reliable estimates across unimodal and bimodal distributions (Ridout and Linkie 2009; Rivero-Monteagudo and Mena 2023). An adjustment factor of one was also used to fine-tune the bandwidth scalar. These parameters were selected to accurately reflect underlying patterns while minimising the impact of noise or over-smoothing (Rivero-Monteagudo and Mena 2023). Overlap coefficients (Δ) were calculated to quantify the extent of diel activity overlap for the same guild between forest and cashew orchards. Three overlap measures— Dhat1 (Δ1), Dhat4 (Δ4), and Dhat5 (Δ5)—were considered. Following Ridout and Linkie (2009), Dhat4 was chosen for its consistent performance across varying sample sizes. As an exception, Dhat 1—best used when at least one of the species (or groups) has fewer than 50 observations—was used for analysing the overlap in the diel activity patterns for both

*Genetta pardina* and *Euxerus erythropus*. The overlap coefficient ranges from 0 (no overlap) to 1 (complete overlap). Confidence intervals at 95% certainty were estimated using the bootstrap method with 9,999 resamples (Ridout and Linkie 2009). These calculations were performed using the ‘overlap’ R package (Meredith and Ridout 2014). To determine whether two activity sets came from identical distributions, we applied a probability test with the ‘compareCkern’ function from the ‘Activity’ R package (Rowcliffe et al. 2014). All analyses were conducted using R software version 4.1.2 (R Core Team 2022).

## RESULTS

Of the 25 species recorded, 13 were nocturnal and 12 were generally diurnal, including two cathemeral species, three diurnal and crepuscular species, and seven strictly diurnal species. Overall, nocturnal species accounted for 81.7% (*N* = 685) of the records and diurnal species for 18.3% (*N* = 153) (Table S2).

In forests, nocturnal species accounted for 82.8% of the mammal records, while diurnal species accounted for 17.2% (Fig. 2a). Here, nocturnal species were mostly represented by *Cricetomys gambianus* (56.4%) and *Ichneumia albicauda* (47.9%), and diurnal species by the squirrels *Euxerus erythropus* (8.2%) and *Funisciurus pyrropus* (6.2%). Similarly, in cashew orchards, nocturnal species accounted for 78.3% and diurnal species for 21.7% (Fig. 2b). Here, nocturnal species were mostly represented by *C. gambianus* (25.7%) and *Genetta pardina* (20.4%) and, to a lesser extent, by *I. albicauda* (16.5%) and *A. paludinosus* (15.0%). Diurnal species were mostly represented by *X. erythropus* (15.3%).

**Figure 2.**
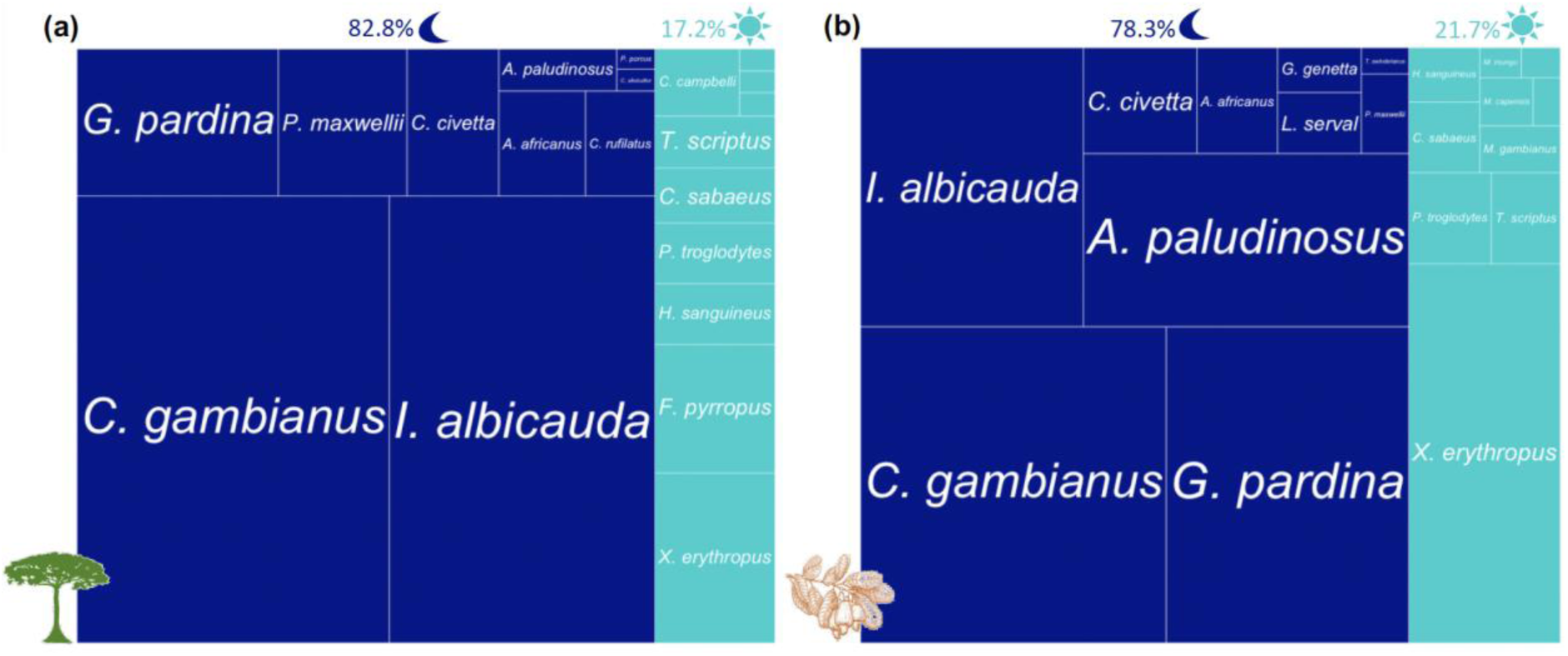
Proportion of the number of records per species per habitat type. (a) forests and (b) cashew orchards. In each panel, each rectangle represents a species, and it is sized according to the proportion of records for the corresponding species. Rectangles are color-coded according to the diel activity guild of each species: nocturnal (dark blue) and diurnal (also including cathemeral species and those with both diurnal and crepuscular activity; light blue). The values provided on the top correspond to the proportion of records for each diel activity guild per habitat type.

Despite the apparent trend for a higher number of records of diurnal mammals in forests (Fig. 3a), the overall activity levels of either diurnal or nocturnal species remained similar between forests and cashew orchards (Table S3, Fig. 3). However, the activity patterns of diurnal species differed between forests and cashew orchards, resulting in less overlap in activity than expected by chance (Δ_4_ = 0.78, *P* = 0.031). In particular, while diurnal activity in the forests peaked in the late morning (∼11h00), that in the cashew orchards peaked in the late afternoon (∼16h30) (Fig. 4a). The activity patterns of nocturnal species remained similar between forests and cashew orchards (Fig. 4b). The same patterns were observed when considering the four most recorded individual species. As such, none of the nocturnal species *C. gambianus*, *I. albicauda* or *G. pardina* varied their activity patterns between forests and cashew orchards (Fig. 5a-c), whereas the activity of *E. erythropus* peaked in the late morning in forests and in the late afternoon in cashew orchards (Fig. 5d).

**Figure 3.**
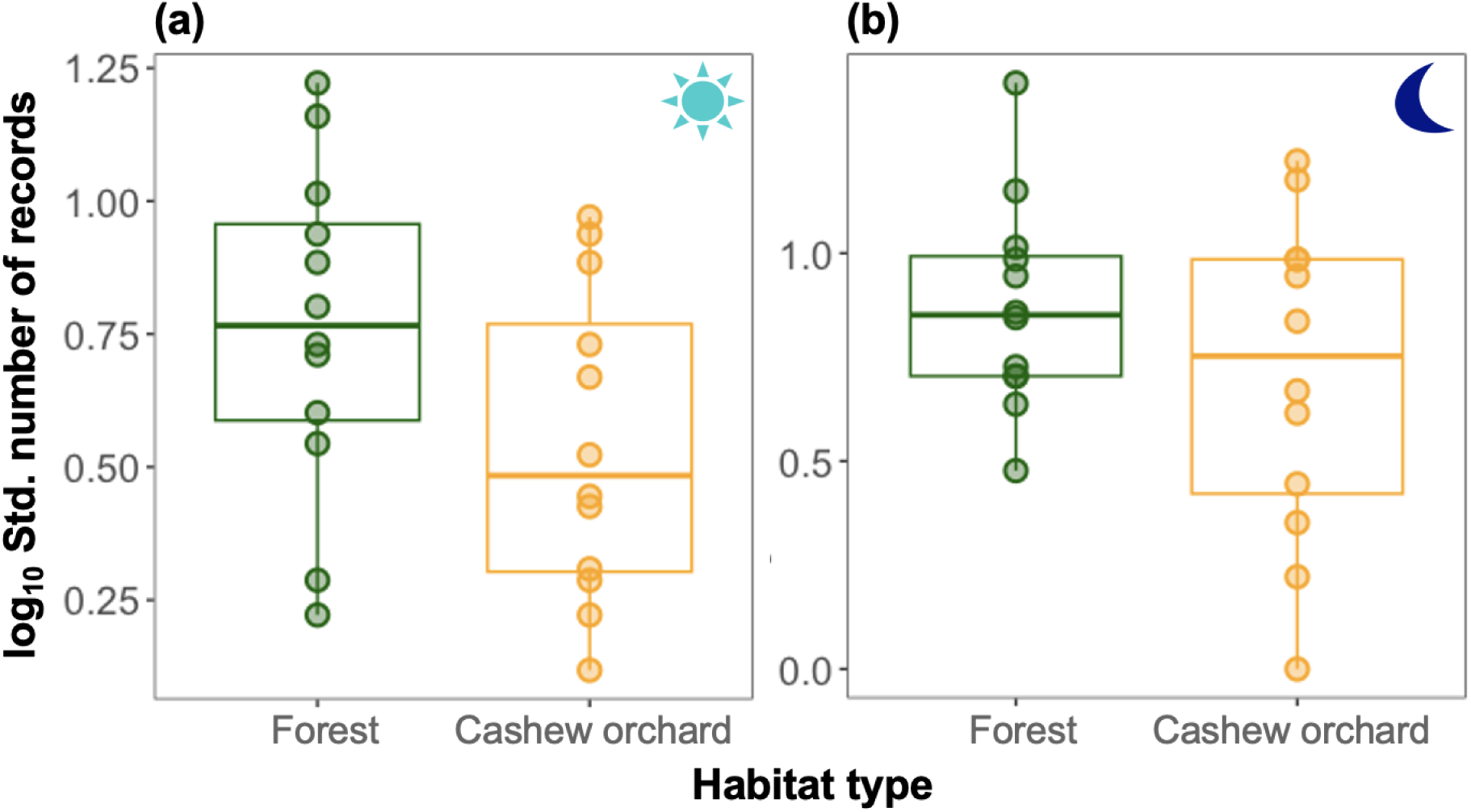
Standardised number of mammal records (log_10_ *x*) per habitat type (forest and cashew orchard) for (a) diurnal mammals (also including cathemeral species and those with both diurnal and crepuscular activity) and (b) nocturnal mammals. Boxplots indicate the median, 1^st^ and 3^rd^ quartiles, and minimum and maximum values.

**Figure 4.**
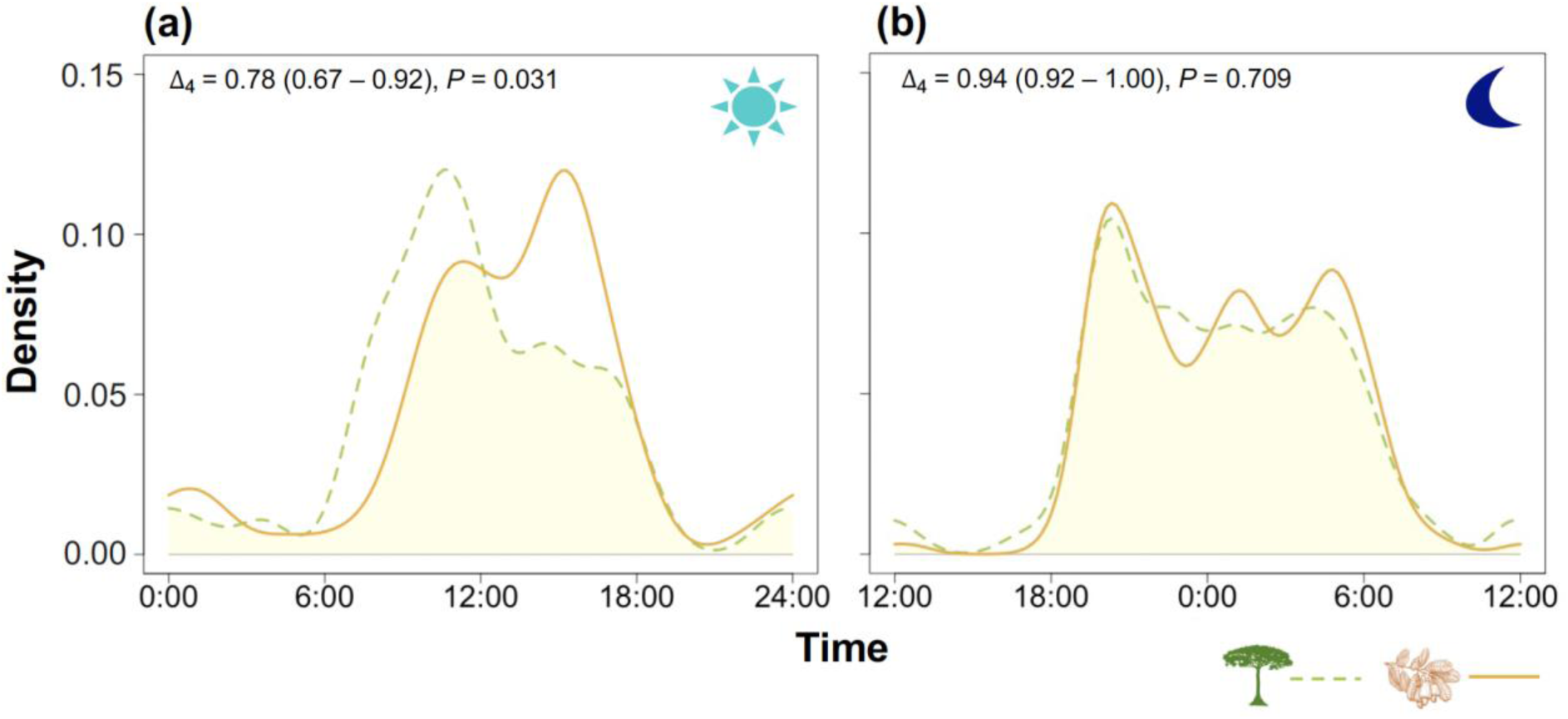
Comparison of the diel activity patterns of (a) diurnal (also including cathemeral species and those with both diurnal and crepuscular activity) and (b) nocturnal mammals co-existing in forests and cashew orchards. For each comparison, the overlap coefficient, and its confidence intervals, as well as the *P*-value (testing whether the probability that the two distributions derive from the same distribution), are provided.

**Figure 5.**
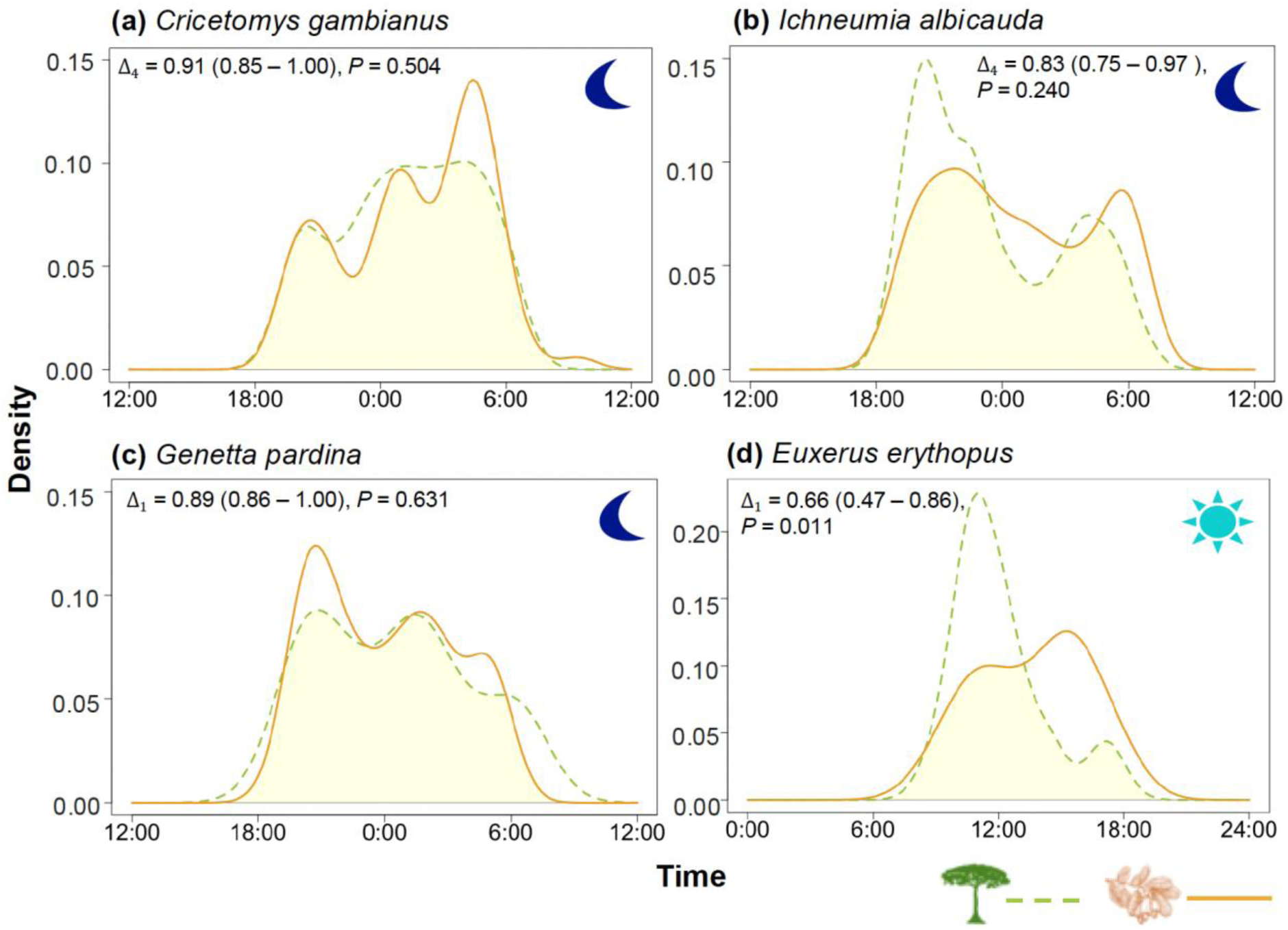
Comparison of the diel activity patterns for each of the four most frequently recorded species (i.e., those with more than 20 detections per habitat type): (a) *Cricetomys gambianus*, (b) *Ichneumia albicauda*, (c) *Genetta pardina*, and (d) *Euxerus erythropus* in forests and cashew orchards. For each comparison, the overlap coefficient, and its confidence intervals, as well as the *P*-value (testing whether the probability that the two distributions derive from the same distribution), are provided. In each panel, the symbols “sun” and “moon” indicate the diel activity guild of the corresponding species (i.e., diurnal, or nocturnal).

## DISCUSSION

Forest conversion to cashew orchards resulted in comparable habitat use by both nocturnal and diurnal mammals. However, the peak of the diel activity of diurnal species shifted from the morning (∼11:00) in the forests to the afternoon (∼16:30) in the cashew orchards. This pattern was also observed when the ground squirrel *E. erythropus*—the most recorded diurnal mammal—was considered separately. In contrast, no significant differences were observed in the diel activity patterns of nocturnal species even when considering the most recorded species separately (i.e., *Cricetomys gambianus*, *Ichneumia albicauda* and *Genetta pardina*). Overall, the shift in the activity peak of diurnal species may reflect behavioural adaptations to the altered habitat.

The temporal partitioning of diel activity (nocturnal vs. diurnal) reduces direct competition for resources like food and space (Bennie et al. 2014). This may allow more species to coexist in human-modified landscapes, with constrained resource availability (Rege et al. 2023), such as cashew orchards. Nocturnal species accounted for 13 out of the 25 (52%) of the mammal species recorded in the Cantanhez National Park, but they amounted to 82.8% and 78.3% of the records in the forests and in the cashew orchards, respectively.

Considering the number of records as a proxy for the species abundance, the higher abundance of nocturnal species is consistent with the expectations for human-modified landscapes, as human disturbance generally favours nocturnality in wildlife (Gaynor et al. 2018a, Burton et al. 2024).

In contrast to forests, cashew orchards represent simplified habitats characterised by greater canopy openness, and lower three height, density, and richness (Nhassé et al. 2024). Cashew orchards are then expected to have lower resource availability and different microclimatic conditions, eventually supporting lower species diversity (Rege et al. 2023; Guedes et al. 2024). Using the same data as here, Na Mone et al. (*under review*) observed that fewer mammal species tended to use cashew habitat, but with varying intensity depending on the trophic guild, with carnivores being more active, and insectivores and herbivores less active in the cashew orchards. In addition to different resource availability and microclimatic conditions, cashew orchards are also subject to some degree of human use during the day (e.g., farm cleaning and cashew nut harvesting; Monteiro et al. 2017; Sierra-Baquero et al. 2024). We would then expect species with diurnal activity to be also less active in the cashew orchards. As mammal activity was similar between forests and cashew orchards regardless of the diel activity guild, it seems that diurnal mammals alternatively cope with the conversion of forests into cashew orchards by delaying the peak of activity.

This change slightly deviates from the increased nocturnality of mammals typically observed in human-disturbed areas (Gaynor et al. 2018a; Li et al. 2021), as despite the delay in the activity, the peak was kept within daytime. Overall, farmers tend to concentrate their activity in the cashew orchards during the morning hours (AFP, pers. obs.), so delaying the peak of the activity may be sufficient for diurnal mammals to avoid direct contact with humans (Gaynor et al. 2018b). To some extent, it is also possible that the microclimate contributes to the observed delay in activity of diurnal species in the cashew orchards. Due to the more open canopy cover, cashew orchards are generally subject to a higher light penetration and thus higher temperatures. The ground squirrel is a small-bodied diurnal species known to have a wide habitat tolerance and is therefore typically observed in agricultural habitats (Cassola 2016). The delayed activity observed in this species may also represent a temperature regulation behaviour, as noted for another species within the same genus (e.g., Scantlebury et al. 2012).

On the contrary, nocturnal species showed consistent activity patterns between forests and cashew orchards. In fact, because human activity in the cashew orchards occurs during the day, nocturnal species do not need to shift their diel activity patterns. A meta-analysis by Gaynor et al. (2018a) examined the effects of human disturbance on mammalian diel activity patterns and found that while many species increased their nocturnality to avoid humans, those that were already nocturnal showed minimal changes in their activity timing. This suggests that nocturnal species are less affected by diurnal human activities and can continue with their natural behaviour without significant temporal adjustments (Procko et al. 2023).

Our findings represent a step towards understanding the behavioural responses of biodiversity to land-use change in the Afrotropics. However, a caveat to our findings is imposed by the limited sampling size, both in terms of sampling sites (*N* = 24) and sampling nights (30 days), resulting in a reduced number of records per species per sampling site. This limitation prevented us from performing species-specific analyses for additional diurnal and cathemeral species. As a delayed activity of diurnal species due to land-use change may have consequences for the prey-predator dynamics (Patterson et al. 2024), we recommend that future studies consider species-level responses, as well as for seasonality.

Overall, our findings highlight that the adaptability of the diurnal mammals in Guinea-Bissau to forest conversion to cashew orchards is mediated by their ability to adjust their diel activity patterns. However, by shifting the peak of their activity from the morning to the afternoon, human disturbance resulted in a “temporal habitat” change (Rivera et al. 2022). Conservation strategies should consider maintaining habitat features that support both nocturnal and diurnal species to ensure ecosystem balance and biodiversity.

## ACKNOWLEDGEMENTS

AFP was supported by the European Union’s Horizon 2020 research and innovation programme under grant agreement No 854248 (TROPIBIO). This programme further funded the fieldwork. We thank: the Institute of Biodiversity and Protected Areas (IBAP) of Guinea-Bissau for authorising and supporting the surveys, namely its director Aissa Regalla and CNP director Queba Quecuta; everyone who provided us support and welcomed us at the accommodation in Iemberém, namely Mamadú Baldé, Joia Tamba, Julmira Na Rescré, Hoina Na Cul, Mamadu Galisa, Djebu Seide, Abubacar Serra, Bubacar Na N’caba, Cadidjatu Galisa, Aissato Camara, Aissatu Turré, and Braima Camara, and all the landowners who allowed us to survey in their cashew farms; the CIBIO-CTM lab staff, namely Susana Lopes and Patrícia Ribeiro, for their support in the laboratory; and Mamadu Cassamá for his invaluable guidance in the field. LP was supported by national funds through FCT – Fundação para a Ciência e a Tecnologia, respectively InBIO Programático FUI 2020-2023 (UIDP/50027/2020).

## AUTHOR CONTRIBUTIONS

AFP conceived the ideas which were improved by comments from all authors; DN, JS, RO, IN, and AFP collected the data; DN and LP processed the camera-trap records; DN analysed the data and led the writing; and all authors contributed with comments and revisions to drafts of the manuscript.

## DATA AVAILABILITY

The data used in this study will be soon available in a data paper (Palmeirim et al., *in prep.*). In the meantime, data will be made available on request.

