## Supplementary Material for "Forest conversion to cashew orchards delays the diel activity of diurnal mammals in West Africa"

**Table S1.** Sampling (trapping) effort per sampling site used to survey mammal assemblages in the Cantanhez National Park, Guinea-Bissau. Site name, geographic coordinates, habitat type, name of the nearest village and number of trap-days (sampling effort) are indicated for each sampling site.

| | Geographic coordinates | | | | |
|---|---|---|---|---|---|
| **Site name** | **North** | **West** | **Habitat type** | **Nearest village** | **No. trap-days** |
| Cam1 | 11.20968 | 15.03850 | Forest | Cambeque | 30 |
| Cam2 | 11.20470 | 15.03853 | Forest | Cambeque | 30 |
| Cam3 | 11.19866 | 15.03882 | Forest | Cambeque | 30 |
| Cam4 | 11.19398 | 15.03956 | Forest | Cambeque | 30 |
| Lau1 | 11.21769 | 15.01977 | Forest | Lauchande | 32 |
| Lau2 | 11.21359 | 15.01813 | Forest | Lauchande | 32 |
| Lau3 | 11.20964 | 15.01949 | Forest | Lauchande | 32 |
| Lau4 | 11.20558 | 15.02135 | Forest | Lauchande | 32 |
| Mad1 | 11.24302 | 15.05483 | Forest | Madina | 30 |
| Mad2 | 11.24527 | 15.05775 | Forest | Madina | 30 |
| Mad3 | 11.25058 | 15.06073 | Forest | Madina | 29 |
| Mad4 | 11.25405 | 15.06513 | Forest | Madina | 30 |
| Cashew1 | 11.25760 | 15.07673 | Cashew orchard | Lauchande | 32 |
| Cashew2 | 11.25923 | 15.08227 | Cashew orchard | Lauchande | 32 |
| Cashew3 | 11.24141 | 15.05224 | Cashew orchard | Lauchande | 24 |
| Cashew4 | 11.24167 | 15.04611 | Cashew orchard | Lauchande | 32 |
| Cashew5 | 11.22479 | 15.01646 | Cashew orchard | Cambeque | 28 |
| Cashew6 | 11.22775 | 15.01821 | Cashew orchard | Cambeque | 30 |
| Cashew7 | 11.22791 | 15.02426 | Cashew orchard | Cambeque | 15 |
| Cashew8 | 11.23022 | 15.02710 | Cashew orchard | Cambeque | 30 |
| Cashew9 | 11.22206 | 15.03895 | Cashew orchard | Madina | 30 |
| Cashew10 | 11.21507 | 15.03737 | Cashew orchard | Madina | 30 |
| Cashew11 | 11.21808 | 15.04491 | Cashew orchard | Madina | 30 |
| Cashew12 | 11.22261 | 15.03015 | Cashew orchard | Madina | 29 |

**Table S2.** Classification of the diel activity patterns, number of records and proportion of records in each land-use type for each species recorded in the Cantanhez National Park, Guinea-Bissau.

| ORDER/ Family | Scientific name | Common name | Activity | No. of records | % Forest | % Cashew |
|---|---|---|---|---|---|---|
| ARTIODACTYLA | | | | | | |
| Bovidae | | | | | | |
| | *Cephalophus rufilatus* | Yellow-backed duiker | Nocturnal | 9 | 100.00 | 0.00 |
| | *Cephalophus silvicultor* | Red-flanked duiker | Nocturnal | 1 | 100.00 | 0.00 |
| | *Philantomba maxwellii* | Maxwell's duiker | Nocturnal | 26 | 96.15 | 3.85 |
| | *Tragelaphus scriptus* | Bushbuck | Diurnal /Crepuscular/ Nocturnal | 12 | 66.67 | 33.33 |
| Suidae | | | | | | |
| | *Potamochoerus porcus* | Red river hog | Nocturnal | 1 | 100 | 0 |
| CARNIVORA | | | | | | |
| Felidae | | | | | | |
| | *Leptailurus serval* | Serval | Nocturnal | 4 | 0.00 | 100.00 |
| Herpestidae | | | | | | |
| | *Atilax paludinosus* | Marsh mongoose | Nocturnal | 50 | 12.00 | 88.00 |
| | *Herpestes ichneumon* | Egyptian mongoose | Diurnal | 2 | 50.00 | 50.00 |
| | *Herpestes sanguineus* | Common slender mongoose | Diurnal /Crepuscular/ Nocturnal | 12 | 75.00 | 25.00 |
| | *Ichneumia albicauda* | White-tailed mongoose | Nocturnal | 196 | 82.14 | 17.86 |
| | *Mungos gambianus* | Banded mongoose | Diurnal/ Crepuscular | 4 | 25.00 | 75.00 |
| | *Mungos mungo* | Gambian mongoose | Diurnal/ Crepuscular | 1 | 0.00 | 100.00 |
| Mustelidae | | | | | | |

| | | | | | |
|---|---|---|---|---|---|
| *Mellivora capensis* | Honey badger | Diurnal | 2 | 0.00 | 100.00 |
| Viverridae | | | | | |
| *Civettictis civetta* | African civet | Nocturnal | 27 | 62.96 | 37.04 |
| *Genetta genetta* | Pardine genet | Nocturnal | 3 | 0.00 | 100.00 |
| *Genetta pardina* | Common genet | Nocturnal | 98 | 41.84 | 58.16 |
| *Genetta* sp. | Genet | Nocturnal | 3 | 100.00 | 0.00 |
| PRIMATA | | | | | |
| Cercopithecidae | | | | | |
| *Cercopithecus campbelli* | Green monkey | Diurnal | 7 | 100.00 | 0.00 |
| *Chlorocebus sabaeus* | Campbell's monkey | Diurnal | 12 | 66.67 | 33.33 |
| Hominidae | | | | | |
| *Pan troglodytes* | Western chimpanzee | Diurnal | 15 | 60.00 | 40.00 |
| RODENTIA | | | | | |
| Sciuridae | | | | | |
| *Funisciurus pyrropus* | Fire-footed rope squirrel | Diurnal/ Crepuscular | 19 | 100.00 | 0.00 |
| *Heliosciurus gambianus* | Gambian sun squirrel | Diurnal | 2 | 50.00 | 50.00 |
| *Xerus erythropus* | Striped ground squirrel | Diurnal | 65 | 38.46 | 61.54 |
| Hystricidae | | | | | |
| *Atherurus africanus* | African brush-tailed porcupine | Nocturnal | 18 | 61.11 | 38.89 |
| Nesomyidae | | | | | |
| *Cricetomys gambianus* | Giant pouched-rat | Nocturnal | 248 | 69.76 | 30.24 |
| Thryonomyidae | | | | | |
| *Thryonornomys swinderianus* | Marsh cane-rat | Nocturnal | 1 | 0.00 | 100.00 |

**Table S3.** Summary of the Generalised Linear Models explaining diurnal and nocturnal mammal activity (log10 x) according to the habitat type (i.e., forest or cashew orchard) across 24 sampling sites in the Cantanhez Nacional Park, Guinea-Bissau.

| Response | Model parameters | Estimate | Std. error | t-value | P-value |
|---|---|---|---|---|---|
| *Diurnal mammal activity (log10 x)* | | | | | |
| | Intercept (forest) | 0.638 | 0.126 | 5.055 | <0.001 |
| | Habitat type (cashew) | -0.306 | 0.179 | -1.716 | 0.100 |
| *Nocturnal mammal activity (log10 x)* | | | | | |
| | Intercept (forest) | 0.871 | 0.094 | 9.242 | <0.001 |
| | Habitat type (cashew) | -0.167 | 0.133 | -1.252 | 0.224 |
